# Validation of DXS as an attractive drug target in mycobacteria

**DOI:** 10.1101/2024.11.04.621930

**Authors:** T Maila, M Nicolaai, R.D Mbau, G.T.M Mashabela

## Abstract

A rapid emergence in the incidences of Tuberculosis (TB) drug resistance undermines efforts to eradicate the disease and strengthens calls for development of new drugs with novel mechanisms of action. In drug discovery, finding an attractive drug target is as important as finding a good drug candidate. Hence more efforts are made to identify, validate and prioritize drug targets in TB drug discovery. Here, using CRISPRi technology, we showed that *dxs1* transcriptional knockdown attenuated growth of both *Mycobacterium smegmatis* and *Mycobacterium tuberculosis* cultures, and the effect was more profound in the latter. Chemical supplementation of the growth medium with 10 μM of isoprenoid pyrophosphates, thiamine and thiamine pyrophosphate failed to rescue growth of *M. smegmatis* cultures, while partial rescue was observed with addition of menatetrenone, a menaquinone derivative with four isoprenyl groups. Similarly, culture growth could not be rescued by the addition of prenol and isoprenol, which suggested the lack of isoprenoid salvage pathway in mycobacteria. Importantly, and in the context of drug discovery, *dxs1* depleted mutants displayed four-fold more sensitive towards a mixture of isoniazid, rifampicin and ethambutol, suggesting that inhibitors of DXS enzyme or other MEP pathway enzymes could potentiate antimycobacterial effect of the first-line TB drugs. Additionally, *dxs1* depletion increased growth retardation of the mutant in acidic pH and under oxidative stress, conditions that are encountered in activated macrophage compartments. Taken together, our results validated DXS as an attractive drug target that should be prioritized for developments on new antitubercular agents.

## 1. Introduction

Tuberculosis (TB), a human disease caused by *Mycobacterium tuberculosis* (*Mtb)*, has again ascended at the top as the deadliest disease by an infectious agent in the world, despite it being described as a curable disease, responsible for 1.3 million deaths and 10.6 million infections in 2022 (WHO, 2023). Perhaps, more concerning is the reported increase in the incidences of multi-drug resistant TB, a complex form of the disease that reduces treatment success rate from 86 % to 63%. It is therefore necessary to intensify research efforts to accelerate the development of more accurate, affordable, shorter and safer regimens for treating TB infection and disease, especially drug-resistant TB (WHO, 2023). A notable limitation of the current TB treatment is increased vulnerability to cross resistance due to a narrow spectrum of molecular targets that are inhibited by the clinical drugs, hence, a call to identify, validate and prioritize novel targets against which new types of antitubercular arsenals should be developed (Capela et al., 2023). Like many bacteria, mycobacteria rely solely on the Non-mevalonate (MEP) pathway for production of isoprenoid precursors, as opposed to the eukaryotes, including humans that rely exclusively on mevalonate pathway, thus making MEP pathway an ideal candidate for drug targeting (Masini et al., 2013).

Isoprenoids constitute one of the largest and structurally diverse groups of natural products with more than 80 000 members and play key metabolic, structural, and regulatory roles in all kingdoms of life (Tarasova et al., 2023). In mycobacteria, isoprenoids play critical roles in cell development, pathogenesis and virulence (Zhao et al., 2013). For example, menaquinone and heme-O/heme-A, the integral parts of electron transport chain, are isoprenoid derivatives (Collins et al., 1977) and so is the decaprenyl phosphate, a hydrophobic chain made of ten isoprenoid units that functions as a carrier of arabinogalactan building blocks that act as encores for mycolic acids (Kaur et al., 2004). Furthermore, isopentenyl tRNA has been associated with increased translational fidelity (Soman & Ram, 2022). In addition, recent studies have implicated 1-tuberculosinyl-adenosine derivatives, a unique *Mtb* diterpenoid derivatives, in promoting pathogen persistence through stimulation of lipid body formation in alveolar macrophages, and the blockage of endocytic compartments acidification (Layre et al., 2014;Buter et al., 2019a; Bedard et al., 2023).

MEP pathway biosynthetic process consists of seven enzyme-catalysed steps that ultimately results in the formation of two universal isoprenoid precursors, isopentenyl pyrophosphate (IPP) and dimethylallyl pyrophosphate (DMAPP) (Eisenreich et al., 2004). The first step, mediated by the deoxyxylulose-5-phosphate synthase (DXS), is the primary rate-limiting step, with the highest flux control coefficient (Wright et al., 2014; Volke et al., 2019; Di et al., 2023) and involves conjugation of pyruvate and glyceraldehyde 3-phosphate. Subsequent decarboxylation of the intermediate to produce DXP in a random sequential reaction catalytic mechanism (Brammer et al., 2011). The recently reported *Mtb* DXS crystal structure was found to have many previously reported common features shared with other previously reported bacterial enzymes, except for a distinct “fork-like” motif involving a highly conserved water molecule in the active site that could be critical for selective inhibition (Gierse et al., 2022).

Since *Mtb* bacteria produce isoprenoid precursors exclusively via MEP pathway, there has been a lot of interest in the pathway enzymes, especially DXS and for two decades, intensive research has been focusing on developing the enzyme active site specific inhibitors (Rohdich et al., 2001; Argyrou & Blanchard, 2004). Despite successful development of potent DXS inhibitors, with some displaying nanomolar inhibition range against the enzyme, no hits/leads have been thud far been validated against *Mtb* as many identified molecules lack whole cell activities, apparently due to inability to permeate mycobacterial cell wall while others suffer poor target selectively. The lack of progress in this regard warrants investigation to validate MEP pathway enzymes, including DXS as viable drug targets. Notwithstanding, essentiality of DXS in mycobacteria has been confirmed and the gene, *dxs1*, is ranked amongst the most vulnerable targets, in which vulnerability relates gene depletion to the corresponding loss of fitness (DeJesus et al., 2017; Bosch et al., 2021). Due to a large number of vulnerable targets identified in *Mtb*, target prioritization is highly recommended, where vulnerability of the newly identified drug targets gene is assessed under different growth environments, including host cell infection models (Mathis et al., 2021; Bosch et al., 2021). Advancements in CRISPRi has been critical in this regards by affording viable mutants deficient in essential genes or enzymes, which are also critical for whole cell screening in search of potential enzyme inhibitors and/or putative synergistic “drugs” for combination therapy without yet having a molecule in hand as it has been shown that depletion mutants can phenocopy antibiotics targeting the mutated protein (Tomasi & Rubin, 2022; Li et al., 2022). Inversely, screening of CRISPRi libraries against existing antibiotics can lead to identification of genes whose depletion hypersensitizes *Mtb* to drugs that are already in clinical use for TB treatment. Here, using CRISPRi, we confirmed the vulnerability of *dxs1* in mycobacteria and demonstrated that growth inhibition emanating from *dxs1* depletion could not be rescued by the addition of MEP pathway metabolites, and also that *dxs1* hypomorphs were hypersensitive to a mixture of the first-line TB antibiotics, low pH and oxidative stress, thus validating DXS an attractive drug target for development of new antitubercular agents.

## 2. Material and Methods

### 2.1. Reagents and chemicals

Difco Middlebrook 7H9 Broth, Difco Middlebrook 7H10 Agar, and BLL Middlebrook enrichment; oleic acid-albumin-dextrose-catalase (OADC) were purchased from Becton Dickinson (BD). Middlebrook OADC enrichment consisted of the following ingredients (approximate formular per 1000 ml): sodium chloride (8,5 g), dextrose (20,0 g), bovine albumin (fraction V) (50,0 g), catalase (0,03 g), oleic Acid (0,6 ml). Glucose, kanamycin sulfate (streptomyces), Tween 80, glycerol, anhydrotetracycline hydrochloride (ATc), hydrochloric acid (HCl), sodium hydroxide (NaOH), dimethyl sulfoxide (DMSO), geranyl pyrophosphate ammonium salt (GPP), isopentenyl pyrophosphate trilithium salt (IPP), 1 Deoxy-D-xylulose-5-phosphate sodium salt (DXP), thiamine hydrochloride (Thh), thiamine pyrophosphate (ThDP), menaquinone (K2), 3-Methyl-2-buten-1-ol (M2B), 3-Methyl-3-buten-1-ol (M3B), clomazone, itaconic acid, nitrosobenzene, diethylenetriamine/nitric oxide adduct (NO), 30% hydrogen peroxide solution (H2O_2_), isoniazid, ethambutol dihydrochloride, rifampicin, pyrazinamide, methanol (MeOH), and acetonitrile (ACN) were all purchased from Sigma Aldrich. pH calibration solutions (4 and 7) were purchased from CRISON, Glass beads (0,1 mm diameter) were purchased from Biospec. The FastRNA Pro Blue kit (containing RNA pro blue solution and lysing matrix B tube containing glass microbeads) was purchased from IEPSA Medical Diagnostic. The Macherey-Nagel Nucleospin RNA plus kit was purchased from Separations company. TURBO DNA-free kit was purchased from Thermo Fisher Scientific and Transcriptor First Strand cDNA Synthesis Kit was purchased from Roche. The oligonucleotides (oligos), BsmBI-v2, and T4 DNA Ligase were purchased from Inqaba Biotec.

### 2.2. Generation of *dxs1* knockdown (*dxs*^*k*D^) strain

The dxs^KD^ strains of *Mtb* H37Rv and *Msm* MC^2^-155 were generated using PLJR965 and PLJR962 plasmids, respectively, as previously described (J. M. Rock et al., 2017). Briefly, two complementary oligonucleotides, encoding a small guide RNA, corresponding to ∼25 – 27 nucleotide bases (Table S1) of the target genes (*dxs1* (Rv2682c) for *Mtb* and *dxs1* (MSMEG_2776) for *Msm*) were designed to contain a BsmBI-v2 restriction site and purchased from Inqaba Biotec as single-stranded oligos. The complementary oligos were then annealed using a PCR machine. The annealed oligos were ligated using T4 DNA ligase into BsmBI-v2 digested plasmids. The CRISPRi constructs were transformed into *E. coli* competent cells, purified and sequenced to confirm insertion of the target genes complementary oligonucleotides before electroporated into *Mtb* H37Rv and *Msm* MC^2^-155. The transformants, selected on 7H10 OADC medium containing Tween 80 and 20 μg/ml kanamycin were grown to the exponential phase, mixed with glycerol to make 33% (v/v) and stored in a freezer at -80 °C until required.

### 2.3. Growth conditions and phenotypic assessment gene knockdown effect

To prepare starter cultures, 300 µl of thawed cells were inoculated into 10 mL of 7H9 broth medium supplemented with 10% OADC, 0.05% Tween 80, 0.2% glycerol, and 20 µg/ml kanamycin. Anhydrotetracycline (ATc) at a concentration of 100 ηg/ml was used to induce the CRISPRi system when required. Unless otherwise mentioned, the cultures were grown at 37 °C in a shaking incubator at 200 rpm (*Msm*) or non-shaking incubator (*Mtb*) to an optical density at 600 nm (OD_600nm_) of 0.4 – 0.6. For *Msm* strains, the starter cultures were inoculated into 20 ml of 7H9 medium supplemented with 0.2% glucose, 0.05% Tween 80, 0.2% glycerol, kanamycin (20 µg/ml), and ATc (100 ηg/ml) (where necessary) in 100 ml flasks to a starting OD_600nm_ of 0.05 and grown in a shaking (200 rpm) incubator at 37°C. Culture growth was monitored by measuring the OD_600_ every 3 hours for the first 15 hours and then once more at 24 hours using a spectrophotometer. For *Mtb* strains, the starter cultures were inoculated into 10 ml of 7H9 medium supplemented with 10% OADC, 0.05% Tween 80, 0.2% glycerol, kanamycin (20 µg/ml), and ATc (100 ηg/ml) (where necessary) in T75 flasks to a starting OD_600nm_ of 0.1. Cultures were incubated at 37 °C in a non-shaking incubator. Culture growth was monitored by measuring OD_600nm_ every day for 7 days. Growth curves were plotted using GraphPad Prism 8 software.

### 2.4. Real-time polymerase chain reaction (RT-qPCR) validation

Starter cultures of *Msm* dxs^KD^ strains, grown as described in section2.3 above (the same other experients), were inoculated into 20 ml of 7H9 medium supplemented with 0.2% glucose, 0.05% Tween 80, 0.2% glycerol, kanamycin (20 µg/ml), and ATc (100 ηg/ml) (where necessary) in 100 ml flasks to a starting OD_600nm_ of 0.1. Cultures were incubated at 37 °C in a shaking incubator at 200 rpm. After 18 hours, 10 ml cultures were transferred into 15 ml centrifuge tubes and centrifuged at 12 000 x g for 10 minutes at 4 °C. Supernatants were discarded and the pellets were resuspended in 2 ml of RNA Pro Blue solution (IEPSA Medical Diagnostic). The suspensions were transferred into a lysing matrix B tube containing glass microbeads and immediately disrupted by ribolyzing (speed = 4 m.s-1, 40 seconds, 3 times, cooling on ice for 1 min between each session) using FastPrep-24 Classic Bead Beating Grinder (MP Biomedicals). The lysates were centrifuged at 12 000 x g for 15 minutes at 4 °C. Cleared lysates (±700 µL) were transferred into 1.5 mL microcentrifuges. The total cellular RNA was extracted using NucleoSpin RNA plus kit (Macherey-Nagel) following the manufacturer’s instructions and the complementary DNA (cDNA) was synthesized using Transcriptor First Strand cDNA Synthesis kit (Roche). Synthesized cDNA was used for the amplification of gene of interest by PCR, using SYBR Green PCR Master Mix (ThermoFisher Scientific), according to the manufacturer’s instructions. Gene-specific primers were designed using the PrimerQuest Tool from IDT, synthesized by Inqaba biotec and were listed in Table 2. The threshold cycle (2^-ΔΔCT^) method was used to calculate changes in gene expression among the strains.

### 2.5. Preparation of IPP and mass spectrometric analysis

Starter cultures of *Msm* dxs^KD^ strains, grown as described above (section 2.3), were inoculated into 20 ml of 7H9 medium supplemented with 0.2% glucose, 0.05% Tween 80, 0.2% glycerol, kanamycin (20 µg/ml), and ATc (100 ηg/ml) in 100 ml flasks to a starting OD_600nm_ of 0.1. Cultures were incubated at 37 °C in a shaking incubator at 200 rpm. At 6- and 18-hour time intervals, samples (10 ml) were collected into 15 ml Falcon tubes from the cultures and centrifuged at 4000 x g for 10 minutes. Supernatants were discarded, and the pellets were resuspended in 1 ml of a cold extraction solvent (methanol: acetonitrile: water (40:40:20) that had been stored at –20°C, vortexed and placed on ice. Suspensions were transferred into a screw-cap microcentrifuge tubes containing glass microbeads and immediately lysed by bead beating on a FastPrep-24 machine using the following settings 4 m/s for 20 s, 4 times with 2 minutes incubation of ice between cycles. The lysates were centrifuged at 12 000 x g for 5 minutes and the supernatants were transferred into centrifuge tubes containing 0.22 µm filter inserts and centrifuged at 12 000 x g for 1 minute. The filtrates were stored in a freezer at –20°C until required. The HPLC-MS/MS analysis of IPP was performed using a high performance liquid chromatograph (Waters Acquity HPLC) connected to a Waters Xevo TQS Micro tandem mass spectrometer (MS/MS). A Waters Acquity HPLC HSS T3 1.8 μm (100 × 2.1 mm) column was used to separate the compounds and placed in a column oven at 50°C. A gradient system was employed using solvent A (water + 0.1% formic acid) and solvent B (acetonitrile + 0.1% formic acid) using an injection volume of 5 μL. The with the flow rate (0.3 ml/min), 0% of solvent B was eluded from 0 to 2 minutes while 0-70% was eluded in 2-5 minutes and 100% of the solvent was eluded in 10 minutes. For the mass spectrometry, the analysis using Waters Xevo TQS Micro in positive mode was conditioned using the cone energy of 15 V and collision energy of 25 eV. The IPP multiple reaction monitoring (MRM) was 245/79 and 245/158.7 while the lowest detection limit (LOD) was 0.0005 mg/L.

### 2.6. pH effect studies

Starter cultures of *Msm* dxs^KD^ strains were inoculated into 20 ml of 7H9 medium supplemented with 0.2% glucose, 0.05% Tween 80, 0.2% glycerol, kanamycin (20 µg/ml), and ATc (100 ηg/ml) in 100 ml flasks to a starting OD_600nm_ of 0.1, which had been adjusted to pH 5.5, pH 6.0, and pH 7.0. Cultures were incubated at 37 °C in a shaking incubator at 200 rpm, and growth was monitored by measuring OD_600nm_ at intervals of 3 h, 6 h, 15 h, and 24 h using a spectrophotometer.

### 2.7. Chemical complementation of dxs^KD^ CRISPRi strain

Starter cultures of *Msm* dxs^KD^ strains were inoculated into 5 ml of 7H9 medium supplemented with 0.2% glucose, 0.05% Tween 80, 0.2% glycerol, kanamycin (20 µg/ml), and either 10 µg/ml of Thh (1 mg/ml), ThDP (1 mg/ml), DXP (1 mg/ml), IPP (1 mg/ml), FPP (1 mg/ml), or GPP (1 mg/ml) in 15 ml centrifuge tubes to a starting OD_600nm_ of 0.1. The cultures were incubated in a shaking incubator (200 rpm) at 37 °C for 6 hours before ATc (100 ηg/ml) was added and were left to grow for further 24 hours under the same conditions. Culture growth was monitored using spectrophotometer (First passage). After 24 hours, cultures were diluted to an OD_600_ of 0.1 in the same growth medium and 5 ml cultures were grown under the same conditions. The growth was monitored for 24 hours using spectrophotometer (Second passage). After another 24 hours, where necessary (OD_600nm_ higher than 0.5), cultures were diluted to an OD_600nm_ of 0.1 in the same growth medium and 5 ml cultures were grown under the same conditions for another 24 hours (Third passage) while cultures with lower than OD_600nm_ of 0.5 were not diluted.

### 2.8. Chemical-genetic interaction study

#### 2.8.1. Single compounds

Starter cultures of *Msm* dxs^KD^ strains were inoculated to OD_600nm_ of 0.1 into 20 ml of 7H9 medium supplemented with 0.2% glucose, 0.05% Tween 80, 0.2% glycerol, kanamycin (20 µg/ml), and where required, ATc (100 ηg/ml) and either of the following compounds: clomazone (320 µg/ml), nitrosobenzene (50 µg/ml), hydrogen peroxide (10 mM) and nitric oxide (1 mM). Cultures were grown in a shaking incubator (200 rpm) at 37 °C and growth was monitored by measuring OD_600nm_ every 3 hours for 15 hours, with a final measurement at 24 hours.

#### 2.8.2. Determination of antibiotics IC50 and dose-response the antibiotic combination

Starter cultures of *Msm* dxs^KD^ strains were inoculated to OD_600nm_ of 0.1 into 20 ml of 7H9 medium supplemented with 0.2% glucose, 0.05% Tween 80, 0.2% glycerol, 20 µg/ml kanamycin and incubated at 37 °C in a shaking incubator until OD_600nm_ of between 0.5 – 0.8. The culture was centrifuged at 5 000 rpm for 10 minutes at room temperature, the supernatant was discarded and the pellet was resuspend in the same medium to achieve OD_600nm_ of 0.1. Stock solutions (1 mg/ml) of solutions of isoniazid, rifampicin and ethambutol, prepared in dimethyl sulfoxide, were diluted with the growth medium to achieve 10 ml each of 80 µg/ml isoniazid, 4.0 µg/ml rifampicin and 2.0 µg/ml ethambutol. The antibiotic solutions were doubly diluted four times by transferring 5 ml into a flask containing 5 ml of the growth medium. The sixth flasks contained 5 ml growth medium alone and were used as negative controls. Into flasks containing 5 ml antibiotic solutions, 5 ml of the diluted culture was added and mixed. The flasks were incubated at 37 °C in a shaking incubators overnight followed by monitoring of culture growth at OD_600nm_, which was analysed GraphPad Prism to determine the antibiotics IC_50_ To prepare for the antibiotic combination dose response, starter culture of *Msm* dxs^KD^ strains was inoculated into 20 ml of 7H9 medium supplemented with 0.02% glucose, 0.05% Tween 80, 0.2% glycerol, and kanamycin (20 µg/ml), in 100 ml flask to a starting OD_600nm_ of 0.1. The culture was grown in a shaking incubator (200 rpm) at 37 °C to an OD_600nm_ of 0.6 – 0.8. The culture was diluted in the same medium containing kanamycin (40 µg/ml) to an OD_600nm_ of 0.2. Antibiotics mixture (5 ml) was prepared in the same medium (ATc was added at 200 ηg/ml where necessary) containing two times IC_50_ concentration of each antibiotic: rifampicin (1.2 µg/ml), isoniazid (37 µg/ml), ethambutol (0.4 µg/ml), 1 ml aliquoted into row 1 on 24 flat-bottom well plate and doubly diluted from rows 1 to rows 2 - 6, which contained 500 µl of the same medium. The diluted cultures (500 µl) were aliquoted into the antibiotics solution in the 24 well plate to make final volumes of 1 ml and the following antibiotic concentrations: rifampicin (0.6 µg/ml), isoniazid (18.5 µg/ml), ethambutol (0.2 µg/ml). The plate was incubated at 37 °C in a shaking incubator (200 rpm). Bacterial growth was monitored by measuring OD_600nm_ after 24 hours.

### 2.9. Isoprenol assimilation

Starter cultures of *Msm* dxs^KD^ strains were inoculated into 20 ml of 7H9 medium supplemented with 0.2% glucose, 0.05% Tween 80, 0.2% glycerol, kanamycin (20 µg/ml), and ATc (100 ηg/ml) (where necessary) and treated with either 3-methyl-2-buten-1-ol (M2B) or 3-methyl-3-buten-1-ol (M3B), or in combination, in 100 ml flasks to a starting OD_600nm_ of 0.1. Cultures were grown in a shaking incubator (200 rpm) at 37 °C, and growth was monitored by measuring OD_600nm_ at intervals of 3 h, 6 h, 9 h, 24 h, and 48 h using a spectrophotometer.

### 2.10. Data and statistical analysis

Data were analysed using GraphPad Prism 8 software and graphs were plotted as means and standard deviations.

## 3. Results and discussion

### 3.1. Addition of ATc reduced growth of *Msm* and *Mtb dxs1* CRISPRi strains

Although *Mtb* contains two putative DXS encoding genes, only *dxs1* (Rv 0111) was found to encode a functional enzyme (Brown et al., 2015) and its essentiality has been demonstrated in many microorganisms that exclusively rely on MEP pathway for the synthesis of isoprenoids, including *Mtb* (DeJesus et al., 2017;Sassetti et al., 2003). The first experiment was to confirm essentiality of *dxs1* in *Mtb* and establish the same in *Msm* using dCas9 CRISPR knockdown system. Accordingly, small guide-RNA sequences were selected down stream of strong PAMs on MSMEG_2776 (*Msm*) and Rv0111 (*Mtb*) genes (see **Table S1**)and cloned into PLJR962 and PLJR965 CRISPRi plasmis, respectively. The CRISPRi plasmids were then electroporated into the respective mycobacterial strains and the CRISPRi systems were activated by addition of 100 ηg/ml ATc. As seen on **Figure 1**, both hypomorphs were restricted for growth, thereby confirming the essentiality of *dxs1* genes for both bacteria.

**Figure 1:**
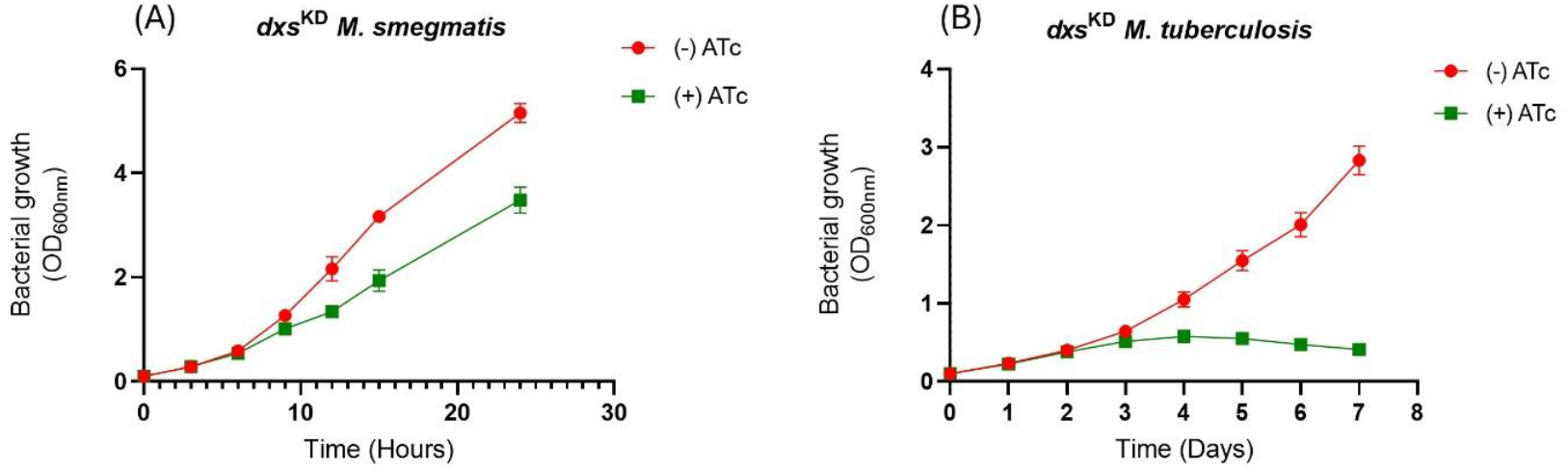
Culture growth of *Msm* (**A**) and *Mtb* (**B**) *dxs*^KD^-CRISPRi strains. The CRISPRi strains were generated by targeting *dxs1* gene with sgRNA presided by strong PAMs. Cultures of *Msm* were grown in 10 ml 7H9 medium supplemented with 0.2% glucose, 0.1% glycerol, 0.05% Tween-80, 50 µg/ml kanamycin in a 37 °C incubator with shaking while *Mtb* cultures were grown in 10 ml 7H9 medium supplemented with 10% OADC, 0.1% glycerol, 0.05% Tween-80, 50 μg/mg kanamycin in a 37 °C incubator without shaking. Intracellular level of *dxs1* RNA transcript was reduced by addition of ATc and corresponded with the reduction of IPP concentration in the cell pellets of *Msm dxs1* CRISPRi strain.

After confirming essentiality of *dxs1* in both strains, subsequent investigations were conducted with *Msm* due to (1) both organisms exclusive reliance on MEP pathway for isoprenoid synthesis (Mancini et al., 2019), (2) high degree of similarity of MEP pathway enzyme homologs between the two organisms, and (see **Table S3**) (Buetow et al., 2007) and (3) for *Msm* rapid’s growth, its consideration as an ideal *Mtb* surrogate that can easily be cultured and ability to be genetically manipulated in BSL-2 laboratory (Sparks et al., 2023). Next, we sought to demonstrate, in three ways, that the observed growth inhibition indeed emanated from d*xs1* depletion. Firstly, transcriptional assessment was conducted and the result demonstrated strong *dxs1* knockdown after 18 hours in ATc treated cultures, clearly confirming on-target knock down (see **Figure 2**A). Secondly, HPLC coupled mass spectrometry (LC-MS/MS) to measure IPP was developed (see **Figure S1**) used to assess the effect of CRISPRi activation, through addition of ATc, on isopentenyl pyrophosphate, the end product of MEP pathway. Here we found that ATc treated cultures had four-fold lower concentrations of IPP compared to the untreated culture (see **Figure 2**B), thereby establishing that a reduction of IPP concentration by as little as four time has a strong fitness cost. This results also highlighted a high demand for a steady supply of isoprenoid buildings for optimal growth in mycobacteria and supported targeting MEP pathway enzymes, including DXS as good drug target (Nahid et al., 2006).

**Figure 2:**
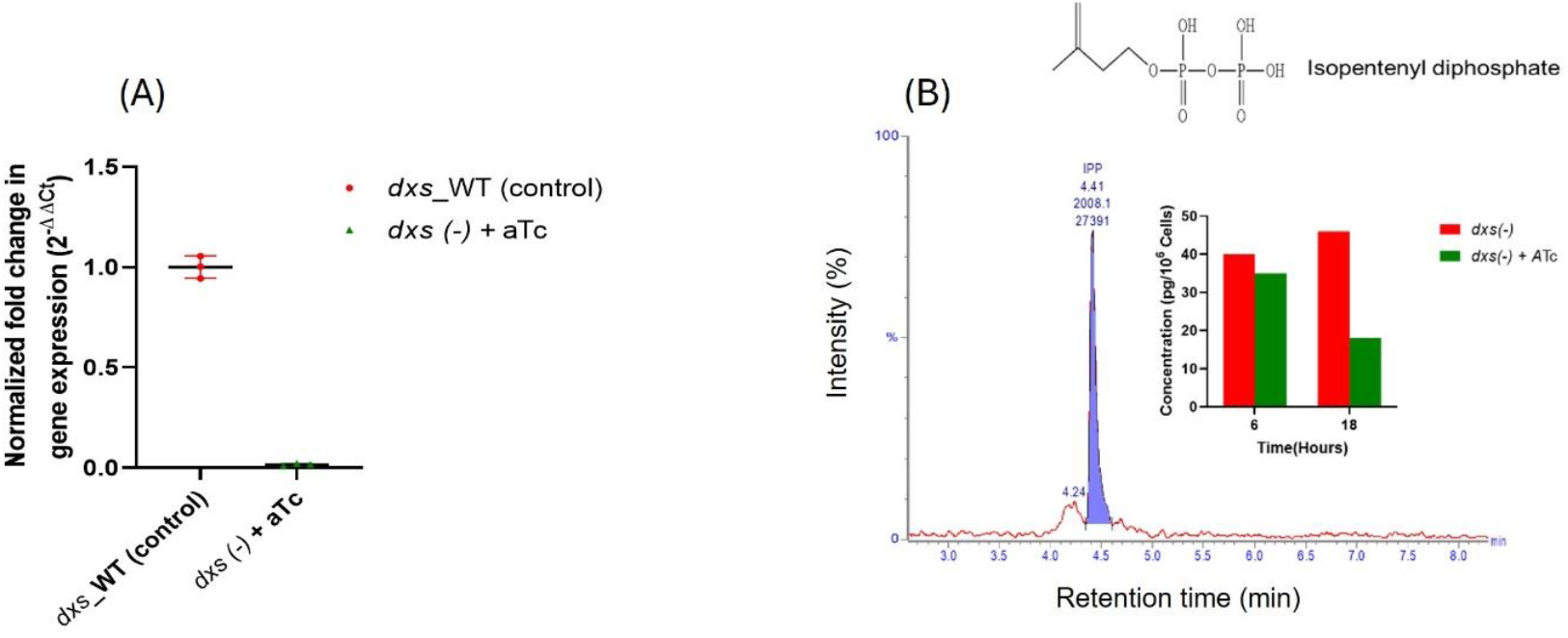
The real time qPCR graph showing transcriptional levels of *dxs1* gene in the cultures of *Msm* CRISPRi strains grown for 18 hours in the medium with and without 100 ηg/ml ATc (**A**) (see **Table S 2** for primers). Mass spectrometric quantification of isopentenyl pyrophosphate isolated from the cell pellets of *Msm* CRISPRi strains grown for 18 hours in the medium with and without 100 ηg/ml ATc (**B**).

### 3.2. Depletion of *dxs1* potentiates inhibitory effect of clomazone, a DXS inhibitor

Finally, the *dxs1* hypomorphs were treated with compounds previously reported to have inhibit DXS enzymatic activity. Clomazone is a well described and highly studied precursor of ketoclomazone, DXS inhibitor (Mueller et al., 2000) while nitrosobenzene disrupts DXS function by competing with glyceraldehyde 3 phosphate as an electron acceptor (Morris et al., 2013). We found that transcriptional knockdown of *dxs1* significantly increased sensitivity to clomazone and nitrosobenzene (see **Figure 3**A and **Figure 3**B). Overall, using whole cell, we provided a significant evidence validating DXS a viable drug target for development of antitubercular agents.

**Figure 3:**
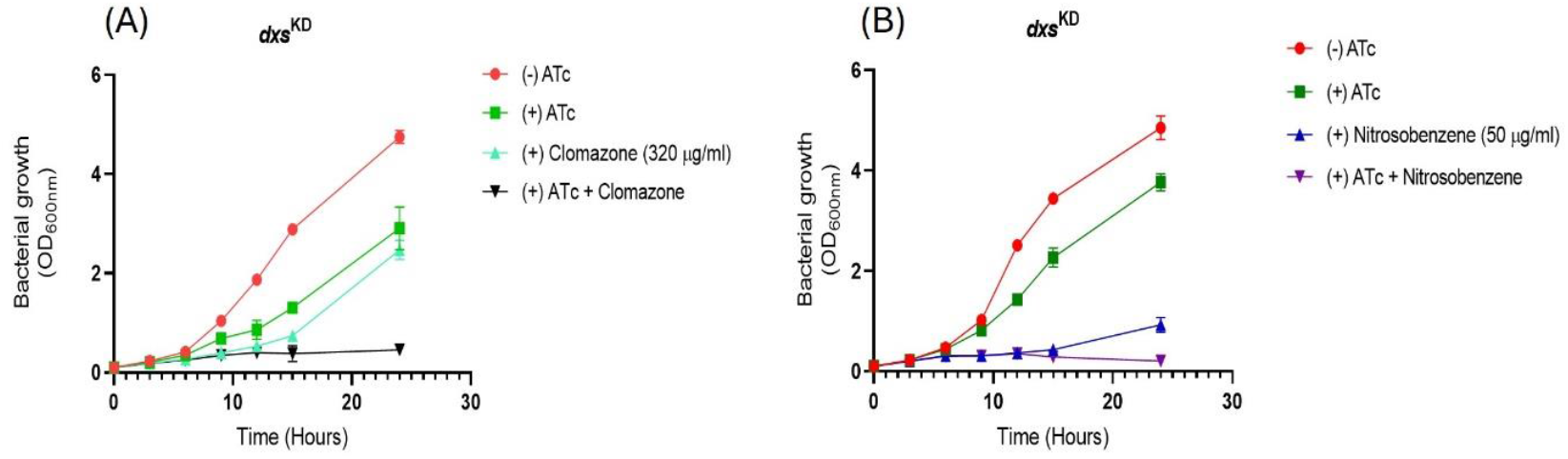
Chemical genetic interaction results of *Msm dxs*1 CRISPRi strain using 320 µg/ml clomazone (A) and 50 µg/ml nitrosobenzene (B). Cultures of *Msm* were grown in 10 ml 7H9 medium supplemented with 0.2% glucose, 0.1% glycerol, 0.05% Tween-80, 50 µg/ml kanamycin and the CRISPRi system was activated by addition 100 ηg/ml ATc.

### 3.3. Growth rescue of *dxs1* hypomorphs

It is highly recommended that target essentiality and vulnerability assessed in different conditions, including those mimicking the human host where a pathogen like *Mtb* reside and cause TB disease (Tomasi & Rubin, 2022). Therefore, we next investigated if *dxs1* depletion-resultant growth inhibition could be off-set by exogenous addition of end products of the MEP pathway and other downstream metabolites as the ability to rescue growth of the mutants could diminish the prospect for DXS as a viable drug target (Woong Park et al., 2011). The rescue experiment involved growing *dxs1* hypomorph for 3 days with a single passage after 24 hour in ATc containing growth medium supplemented with isoprenoid precursor and products, isopentenyl phosphate, geranyl pyrophosphate, farnesyl pyrophosphate, menatetrenone, thiamine and thiamine pyrophosphate. None of the compounds rescued growth of *dxs1* hypomorph at 24- and 48-hour time points (see **Figure 4**A). In day 3, the cultures did show some growth and menatetrenone partially rescued growth of the hypomorph, up to a third of the *dxs1* sufficient, suggesting that *dxs1* depletion, in part, caused menaquinone deficiency. In contrast, no significant growth rescue was observed upon addition of 10 µM of the other supplements, including IPP, which could be attributed to lack of mycomembrane permeability due to the presence of phosphate groups. Failure to rescue *dxs1* hypomorph growth by 10 µM IPP was particularly interesting as IPP levels, produced via mevalonate pathway, are estimated to be between 10 and 100 nM and suggested that host cellular IPP may fail to rescue DXS inhibition *in vivo* (Riganti et al., 2012).

**Figure 4:**
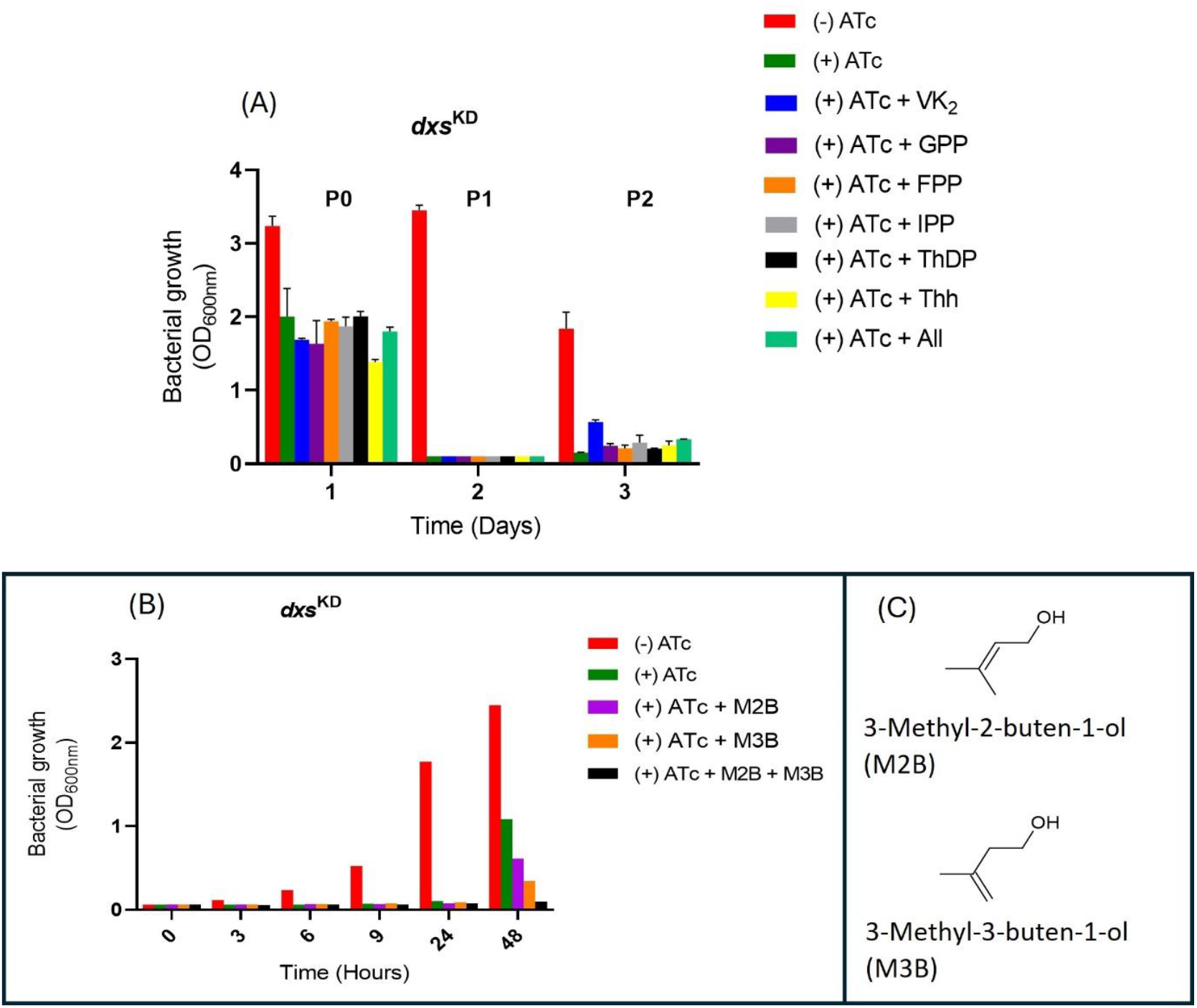
Growth rescue of *dxs1* hypomorph by addition of 10 µM each of isopentenyl phosphate, geranyl pyrophosphate, farnesyl pyrophosphate, menatetrenone, thiamine and thiamine pyrophosphate (**A**). Growth rescue of *dxs1* hypomorph by addition of 5 mM each of 3-methyl-2-buten-1-ol (M2B) and 3-methyl-3-buten-l-ol (M3B) (**B**). Cultures of *Msm* were grown in 10 ml 7H9 medium supplemented with 0.2% glucose, 0.1% glycerol, 0.05% Tween-80, 50 µg/ml kanamycin and the CRISPRi system was activated by addition 100 ηg/ml ATc. Name and chemical structure of the prenol and isoprenol (**C**).

Some bacteria are known to tolerate high concentrations of isoprenol, even up to 25 mg/L without adverse effect on growth (Ferraz et al., 2021), suggesting existence of adaptation mechanisms such as isoprenoid salvage etc. Isoprenoid salvage, a two-step pathway that converts prenols and isoprenols to their respective pyrophosphate occurs in some bacteria, plants, and animals, although it is yet to be described in mycobacteria, (Voshall et al., 2024;Crispim et al., 2024;Lis & Kuramitsu, 2003), despite a report of the existence of isopentenyl phosphate kinase in all three domains of life (Dellas et al., 2013). Therefore, we investigated possible MEP pathway bypass by prenol/isoprenol assimilation in mycobacteria by isoprenol supplementation of the growth medium. Our results showed that supplementation of growth medium with prenol, M2B, and isoprenol, M3B, not only failed to rescue growth, but inhibited the hypomorph growth in an concentration dependent manner (see **Figure 4**B), a phenomenon that was previously also reported in *E. coli* (Ferraz et al., 2021). The observed lack of growth rescue and accompanying toxicity of the isoprenol suggested a possible lack of the necessary kinases for the isoprenoid salvage pathway in mycobacteria. Overall, our results provided evidence that further supported DXS as a good drug target, whose inhibition would not be rescued by MEP pathway intermediates or isoprenoid salvage pathway (Yeh & DeRisi, 2011).

### 3.4. Dxs hypomorphs are hypersensitive to a mixture of first-line TB antibiotics

New multidrug therapies are needed to improve TB treatment outcomes and should include drugs that shorten treatment and increase efficacy, or both (WHO, 2023) Combination therapy holds a promise towards shortening of TB treatment time from the current standard of care of 6 months (Aldridge et al., 2021; Ramón-García et al., 2011). We reasoned that a possible answer to whether new and/or more potent TB treatment can be achieved can also be found in improving effectiveness of the existing combinations by addition of new agents. Drug screening using CRISPRi mutants provide opportunities to predict combination chemotherapy through chemical-genetic interactions where compounds are screened at sub-MIC concentrations to identify genes whose depletion potentiate activity of antibiotic (J. Rock, 2019). Therefore, we sought to investigate whether depletion of *dxs1* would potentiate antitubercular activity of sub-lethal concentrations of antibiotic mixture comprising three first-line TB antibiotics, isoniazid, rifampicin and ethambutol. Pyrazinamide was excluded due to the lack of antimicrobial effect at physiological pH, even at high concentration (data not presented), as previously observed in other studies (Zhang et al., 1999). Prior to constituting the combination mixture, IC_50_ of each antibiotic was determined separately using *Msm* wild type (see **Figure 5**A). In agreement with previous studies, we observed high and moderate susceptibility of *Msm* to ethambutol and isoniazid, respectively (Lelovic et al., 2020; Sakiyama et al., 2023). A mixture was constituted using the corresponding IC_50_ concentration of each antibiotic (isoniazid (15 μg/ml), ethambutol (0.19 μg/ml) and rifampicin (0.75 μg/ml) and used for the chemical-genetic interaction to establish possible potentiation. The results showed that, *dxs1* hypomorph was significantly more susceptible to the antibiotic combination, displaying four-fold more sensitivity (see **Figure 5**B). It has been reported that a mere 2-fold decrease in minimum inhibitory concentration can predict higher rates of TB relapse and resistance (Colangeli et al., 2018). Therefore our results predicted that selective DXS inhibitors will significantly improve effectiveness of the first-line TB treatment.

**Figure 5:**
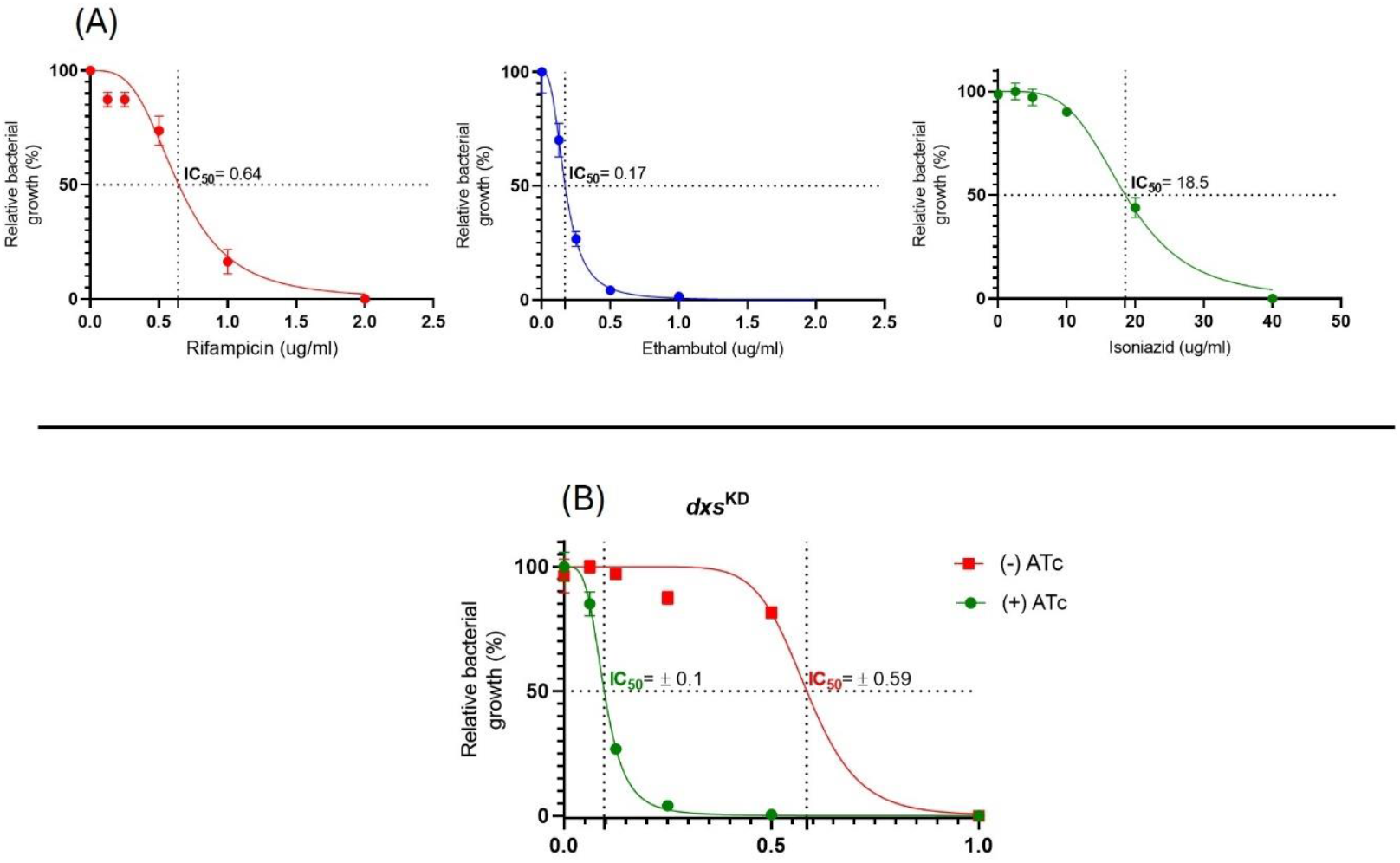
Dose response curve of *dxs1* hypomorphs to increasing concentration of rifampicin, ethambutol, isoniazid (**A**) and a mixture of rifampicin, isoniazid and ethambutol (0.6 µg/ml: 0.2 µg/ml: 18.5 µg/ml) (**B**). Cultures of *Msm* were grown in 10 ml 7H9 medium supplemented with 0.2% glucose, 0.2% glycerol, 0.05% Tween-80, 50 µg/mg kanamycin and the CRISPRi system was activated by addition 100 ηg/ml ATc.

### 3.5. Dxs hypomorphs are hypersensitive to acidic pHs

Survival of *Mtb* in acidic microenvironments in the host depends on the ability to adapt and maintain their acid-base homeostasis and new TB drugs need to be effective under acidic environment (Darby et al., 2013a). Indeed, the decision for the inclusion of pyrazinamide into TB treatment was informed by potentiation of the treatment regimen *in vivo*, which resulted in the shortening TB treatment time (Service, 1979). Identification and targeting of genes products that mediate pH adaptation are highly desirable in the efforts to eradicate recalcitrant *Mtb* pathogens residing in the acidic phagosomal compartments, which varies between pH 4.5 and 8.8 (Darby et al., 2013b;Early, Ollinger, et al., 2019a;Early, Mullen, et al., 2019;Greenstein & Aldridge, 2022). In this regard, we next investigated whether *dxs1* depletion would increase the mutant vulnerability to acid environment; pH 5.5 was chosen because lower pH severely abrogated growth (Vandal et al., 2009a; Chapman & Bernard, 1962; Portaels & Pattyn, 1982; Early, Ollinger, et al., 2019b). Our result showed general pH-dependent growth inhibition from pH 7.0 to 5.5 in both *dxs1* deficient and proficient strains. Depletion of *dxs1* gene did not change the mutant sensitivity to acidic conditions from pH 7 to pH 6. In contrast, *dxs1* depletion increased mutant acid sensitivity from 6.0 to 5.5 (see **Figure 6**A and **Figure 6**B). The increased acid sensitivity was in accord with some previous reports in which MEP pathway associated enzymes and metabolites were found to play roles in the regulation and neutralization of macrophage phagosomal acid (Vandal et al., 2009b). For examples, 1-tuberculosinyladenosine, a cyclic derivative of geranylgeranyl pyrophosphate, which is highly prevalent among patient-derived *Mtb* strains was found to selectively accumulates in macrophage acidic compartments, where it neutralizes the pH and swells lysosomes, destroying their multilamellar structure (Buter et al., 2019b). Additionally, transposon disrupted Rv2136c, a gene that encode for an undecaprenyl pyrophosphate phosphatase, rendered the mutant highly acid sensitive (Vandal et al., 2008). Taken together, our results implicated DXS or MEP pathway metabolites in the regulation of pH in mycobacteria and underscore enzymes of the pathway including DXS as attractive drug targets.

**Figure 6:**
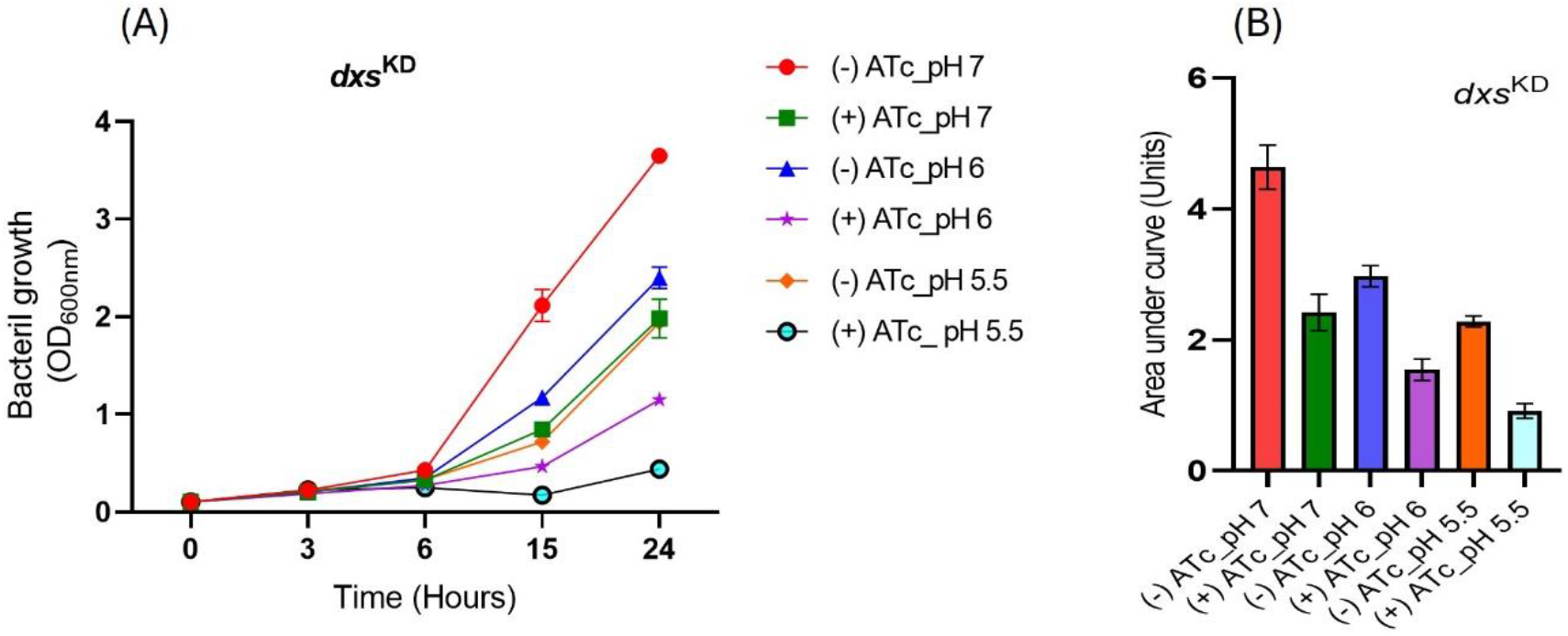
Effect of low pH on *dxs1* CRISPRi strain demonstrated by a linear plot (**A**) and the corresponding area under curve plot (**B**). Cultures of *Msm* were grown at pH 7.0, 6.0 and 5.5 in 10 ml 7H9 medium supplemented with 0.2% glucose, 0.1% glycerol, 0.05% Tween-80, 50 µg/ml kanamycin and the CRISPRi system was activated by addition 100 ηg/ml ATc.

### 3.6. Dxs hypomorphs are hypersensitive to oxidative stress

Production of oxidant is a hallmark of activated macrophages, with reactive oxygen and nitrogen species as the most produced oxidants. Despite its controversy, a highly debatable thesis that all bactericidal antibiotics kill bacteria by generating reactive oxygen or nitrogen species (ROS/RNS) highlight the significance of oxidative stress in antimicrobial drug discovery, thus providing grounds for decades of TB drug discovery research focused on disrupting the pathogen antioxidant systems (Yano et al., 2011; Singh et al., 2008). It was against that background that we sought to investigate potential potentiation of oxidative stress mediated bacterial killing by intracellular reduction of *dxs1* gene. Indeed our results showed increased sensitivity to nitric oxide by the *dxs1* knockdown (see **Figure 7**A), while hydrogen peroxide sensitivity was as moderate, yet not antagonistic (see **Figure 7**B). The increased sensitivity to the oxidative stress could be explained by the presence of two iron sulfur cluster dependent enzymes that mediate the last two steps of MEP pathway, which are highly vulnerable to oxidative stress, especially nitrosative stress (Artsatbanov et al., 2012). Therefore, our results suggested that inhibitors of DXS would work synergistically with oxidants such as those produced by the host *in vivo*, once more supporting prioritization of DXS as an attractive drug target.

**Figure 7:**
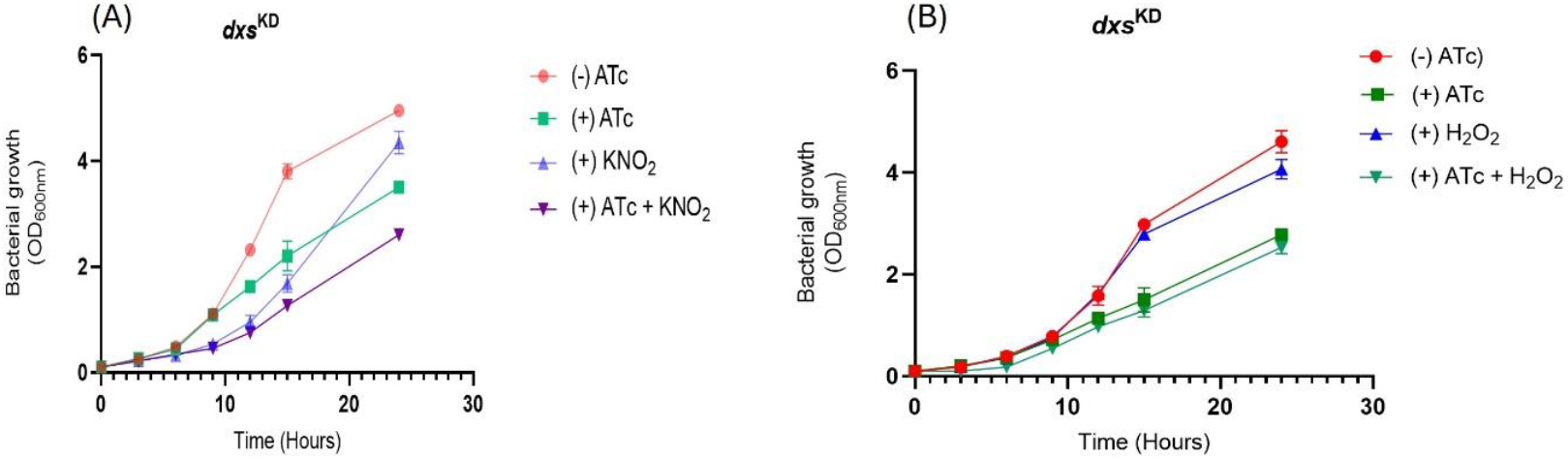
Effect of nitric oxide (1 mM) (**A**) and hydrogen peroxide (10 mM) (**B**) on *dxs1* CRISPRi strain. Cultures of *Msm* were grown in 10 ml 7H9 medium supplemented with 0.2% glucose, 0.1% glycerol, 0.05% Tween-80, 50 µg/ml kanamycin and 1 mM nitric oxide or 10 mM hydrogen peroxide, and the CRISPRi system was activated by addition 100 ηg/ml ATc.

## 4. Conclusion

For successful validation and prioritization of hit drug targets, it is highly recommended to conduct *in vitro* assessment of essential and promising drug target under physiologically relevant conditions to avoid unnecessary and costly drug discovery exercises (Pethe et al., 2010; Sun et al., 2022). Here, we demonstrated that intracellular depletion of *dxs1* led to severe growth impairment in *Msm* and *Mtb*, which was accompanied by over 98% *dxs1* knockdown in *Msm*. Unlike in malaria parasite and spinach. (Yeh & DeRisi, 2011a; Ferhatoglu & Barrett, 2006), we also showed that IPP addition, even at 10 times more than the physiological levels in human, could not rescued growth of *dxs1* hypomorphs. Consistently, *dxs1* depletion led to severe disruption of MEP pathway and the hypomorph were hypersensitive to DXS inhibitors, clomazone and nitrobenzene. The later was particularly important as it confirmed, for the first time, permeability of clomazone through mycobacterial cell wall. Importantly, perhaps more relevant towards TB treatment, *dxs1* depletion increased sensitivity to a mixture of the first-line TB antibiotics by more than four-fold, thus predictive of a positive synergy between DXS specific inhibitors and TB antibiotics, which would impact positively towards reducing TB treatment time. Additionally, our results implicated DXS or MEP pathway in the adaptation to acidic condition and nitrosative stress, the stressors imposed to invading pathogens by activated alveolar macrophages. Taken together, our results elevated DXS as one of important drug targets worthy of prioritization for development of new antitubercular agents, and more work is currently underway in our research group to fully characterize *dxs1* hypomorphs in *Mtb* and search for new inhibitors of DXS.

## 5. Acknowledgements

The authors are grateful for the financial support received from South African Medical Research Council, Grand Challenges Africa Drug Discovery cohort 2, National Research Foundation and Stellenbosch University.

## 7. Supplementary information

**Table S1:**
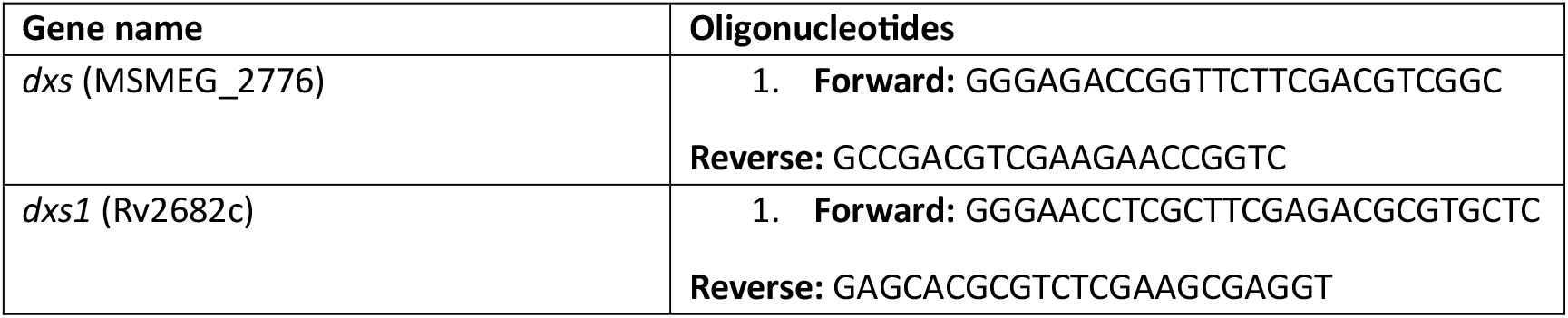
Oligonucleotides used to generate CRISPRi strains.

**Table S 2:**
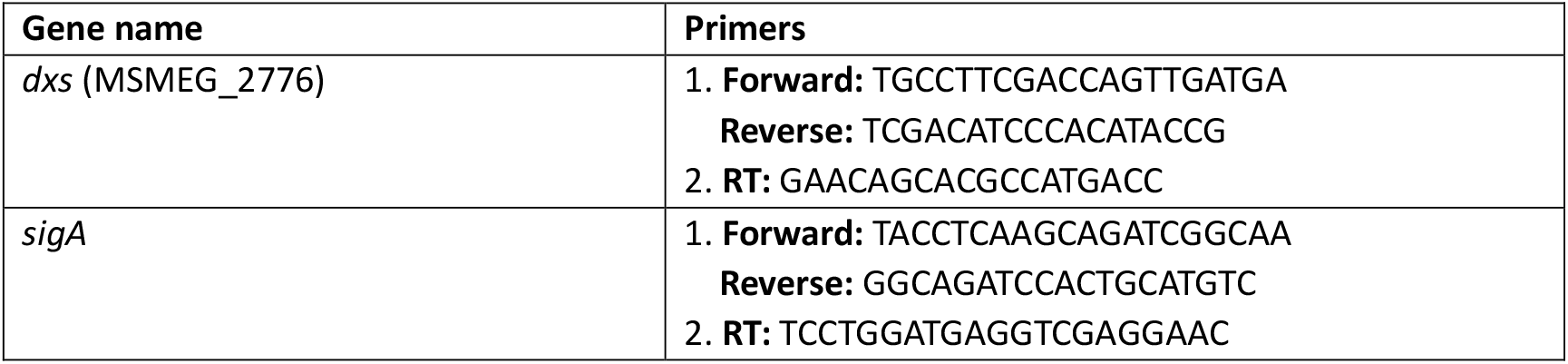
The RT-qPCR primers designed and used in this study.

**Table S3:** Table showing similarities of MEP pathway enzymes in Mycobacterium smegmatis and Mycobacterium tuberculosis.

| Gene |  |  |  |  |  |
| --- | --- | --- | --- | --- | --- |
| Protein | *M. smegmatis* | *M. tuberculosis* | Accession | E-value | Identity (%) |
| DXS | MSMEG_2776 | Rv2682c | WP_011728575. | 0.0 | 79.47 |
| IspC | MSMEG_2578 | Rv2870c | WP_011728448.1 | 0.0 | 77.98 |
| IspD | MSMEG_6076 | Rv3582c | WP_011731019.1 | 2,00E-85 | 62.61 |
| IspE | MSMEG5436 | Rv1011 | WP_045756603.1 | 1E-167 | 79.28 |
| IspF | MSMEG_6075 | Rv3581c | WP_011731018.1 | 1,00E-74 | 75.69 |
| IspG/GcpE | MSMEG_2580 | Rv2868c | WP_036453060.1 | 0.0 | 91.08 |
| IspH/LytB2 | MSMEG5224 | Rv1110 | WP_014878272.1 | 0.0 | 85.71 |

**Figure S1:**
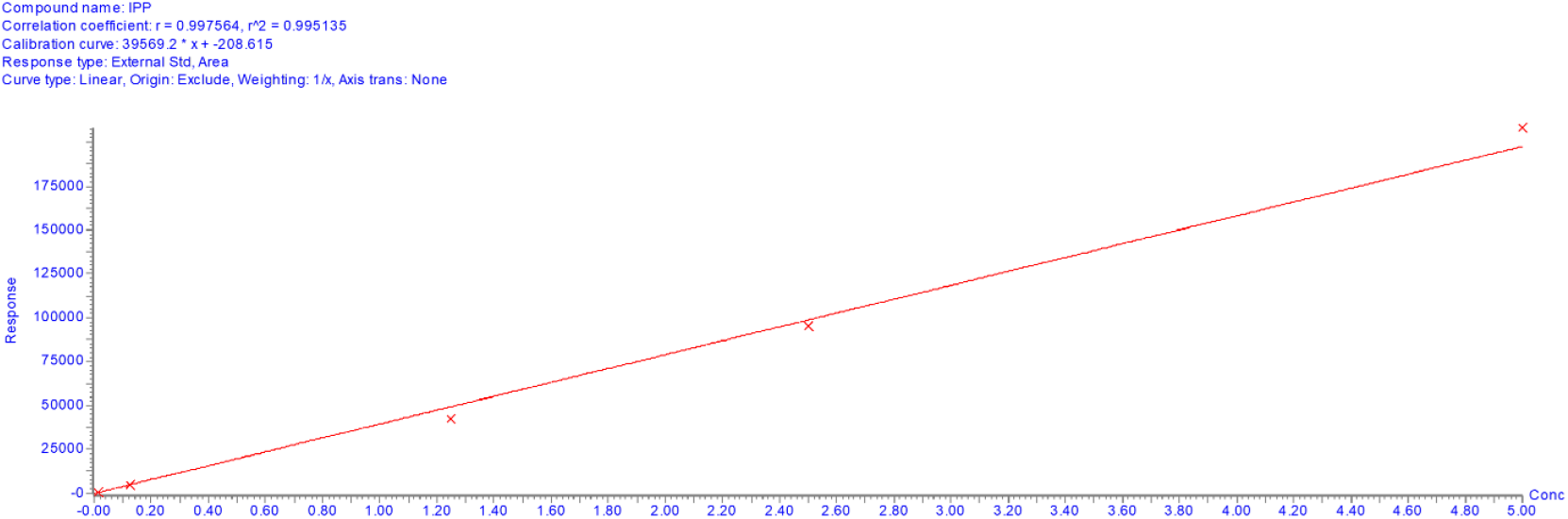
Standard curve of IPP generated using LC/MS/MS mass spectrometry.

